# Supervised classification enables rapid annotation of cell atlases

**DOI:** 10.1101/538652

**Authors:** Hannah A. Pliner, Jay Shendure, Cole Trapnell

## Abstract

Single cell technologies for profiling tissues or even entire organisms are rapidly being adopted. However, the manual process by which cell types are typically annotated in the resulting data is labor-intensive and increasingly rate-limiting for the field. Here we describe Garnett, an algorithm and accompanying software for rapidly annotating cell types in scRNA-seq and scATAC-seq datasets, based on an interpretable, hierarchical markup language of cell type-specific genes. Garnett successfully classifies cell types in tissue and whole organism datasets, as well as across species.

## MAIN

Single-cell transcriptional profiling (scRNA-seq) has emerged as a powerful means of cataloging the myriad cell types present in complex animal tissues (methods reviewed in ref ^1^). The computational steps of constructing a cell atlas typically involves unsupervised clustering of cells based on their gene expression profiles, followed by the annotation of known cell types amongst the resulting clusters^2,3^. With respect to the latter task, there are at least four challenges that are proving rate-limiting for the field. First, cell type annotation is labor intensive, requiring extensive literature review of cluster-specific genes^4^. Second, any revision to the analysis (*e.g.* additional data, adjustment of parameters) necessitates manual reevaluation of all previous annotations. Third, cell type annotations are not easily transferred between datasets generated by independent groups on related tissues, resulting in wasteful repetition of effort. Finally, cell type annotations are typically *ad hoc*; although ontologies of cell types exist^5,6^, we lack tools for systematically applying these ontologies to annotate new scRNA-seq datasets. Collectively, these challenges are strongly hindering progress towards a consensus framework for cell types and the features that define them.

Towards addressing these challenges, we devised Garnett (**Figure 1A**). Garnett consists of four components. First, Garnett defines a markup language for specifying cell types using the genes that they specifically express. The markup language is hierarchical in that a cell type can have subtypes (*e.g.* CD4+ and CD8+ are subsets of T cells). Second, Garnett includes a parser that processes the markup file together with a single-cell dataset, identifying representative cells bearing markers that unambiguously identify them as one of the cell types defined in the file. Third, Garnett trains a classifier that recognizes additional cells as belonging to each cell type based on their similarity to representative cells, similar to an approach that our groups recently developed for annotating a single cell mouse atlas of chromatin accessibility^8^. Importantly, Garnett does not require that cells be organized into clusters, but it can optionally extend classifications to additional cells using either its own internal clustering routines or those of other tools, such as Monocle^9^ or Seurat^10^. Finally, Garnett provides a method for applying a markup file together with a classifier trained on one dataset to rapidly annotate additional datasets.

**Figure 1.**
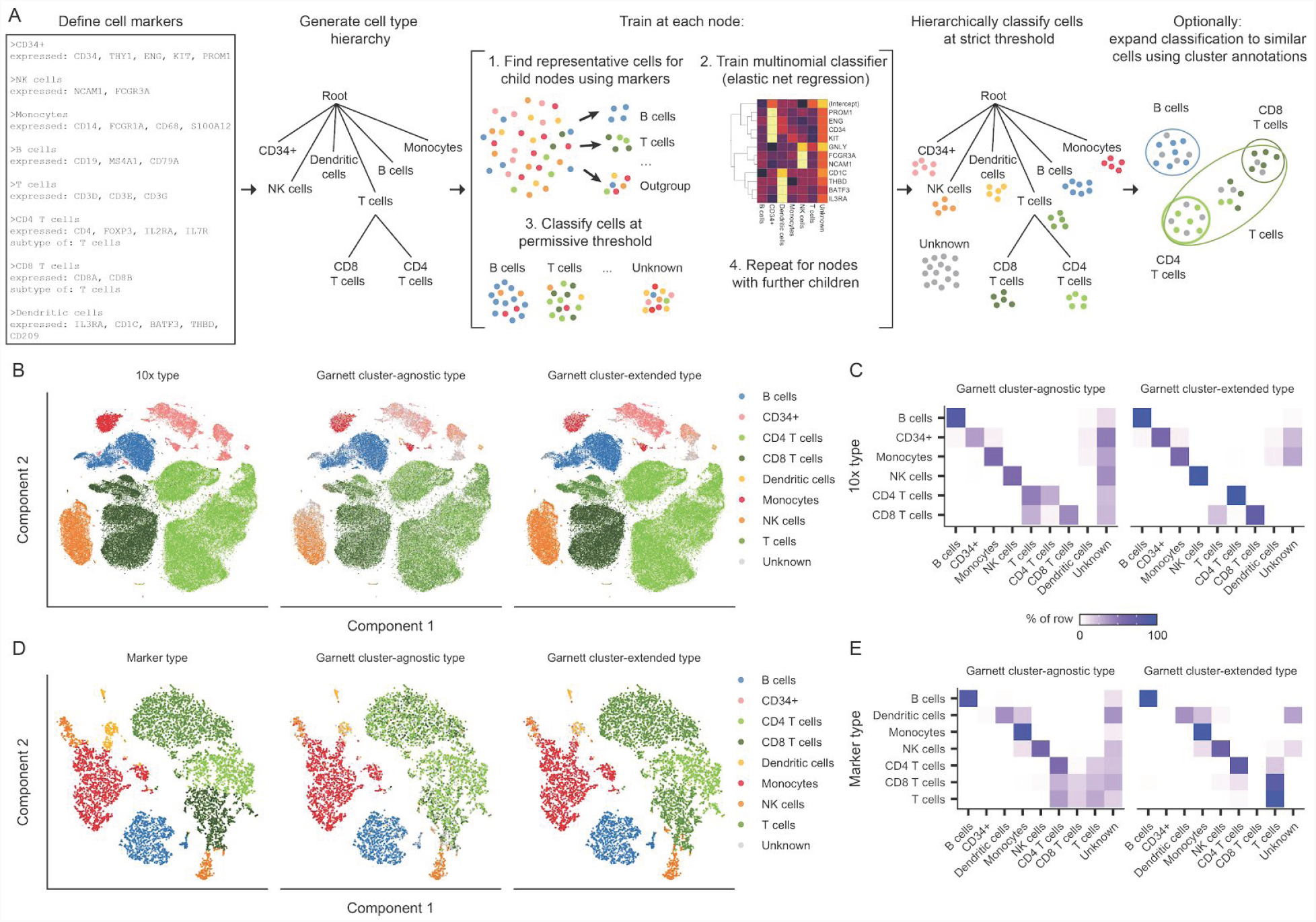
Garnett accurately classifies peripheral blood mononuclear cells. A) Overview of Garnett algorithm. See Methods for algorithmic details. Briefly, Garnett takes as input a marker file that defines cell types using marker genes, and builds a cell type hierarchy that can include cell subtypes. Next, Garnett trains a classifier using elastic net multinomial regression^7^ at each node beginning at the root of the tree by comparing cell type representative cells. Lastly, Garnett hierarchically classifies all cells and optionally provides a second cluster-extended classification. **B)** t-SNE plots of 10x Genomics’ 100,000 cell PBMC dataset. The first panel is colored by cell type based on FACS sorting, the second panel is colored by cluster-agnostic cell type according to Garnett classification, and the third panel is colored by the Garnett cluster-extended type, which labels cells based on the composition of their cluster or community. **C)** A heatmap of data in (B) comparing the labels based on FACS (rows) with the cluster-agnostic (left) and cluster-extended (right) cell type assignments by Garnett (columns). Color represents the percent of cells of a certain FACS type labeled each type by Garnett. **D)** t-SNE plots of 10x Genomics V2 chemistry applied to 8,000 PBMCs from a healthy donor. The first panel is colored by type determined manually using known gene markers. The second and third panels are colored by Garnett cluster-agnostic and cluster-extended cell type assignments by a classifier trained on the data shown in panels (B) and (C). **E)** Similar to panel (C), a heatmap of data in (D).

We tested Garnett on a benchmark single-cell RNA-seq dataset generated using the 10X Chromium platform. This dataset is comprised of 94,571 peripheral blood mononuclear cells (PBMCs) that were individually immunophenotyped via flow cytometry and therefore have a “gold standard” annotation of cell type^11^. Garnett requires at least one marker gene for each cell type, ideally one that is specifically expressed and readily detectable only in that cell type. As a supervised method, Garnett’s accuracy will be dependent on these markers, so we devised a measure of each marker’s usefulness for the purposes of Garnett classification. This marker score combines the number of cells that a marker nominates with an estimate of how many cells inclusion of the marker will render ambiguous. To classify the PBMCs, we populated a marker file including each of the expected cell types using commonly used markers in the literature. We then used Garnett’s quality metric to exclude poorly scoring markers (ambiguity > 0.5) before proceeding with classification (**Supplementary Figure 1A**).

Garnett assigned 71% (3% incorrect, 26% unclassified) of the cells to the correct type (“cluster-agnostic type”), with 34% of T cells also receiving a correct subtype classification (41% not subclassified, 23% unclassified, 2% incorrect) (**Figure 1B-C**). Cells that remained unlabeled were comparably distributed amongst immunophenotypes, suggesting that the algorithm was not failing to recognize one or more of the cell types entirely. Moreover, by expanding cell type assignments to nearby cells using Louvain clustering^12^ (“cluster-extended type”), correct assignments increased to 94% (2% incorrect, 4% unclassified), with 91% of T cells also receiving a correct subtype classification (8% not subclassified, <1% unclassified, <1% incorrect).

We next evaluated Garnett’s ability to classify data not seen during training by analyzing PBMCs that were generated with a second generation of the Chromium system (“V2”). These cells were unsorted and profiled using a different library preparation method that yields much greater molecular depth per cell (**Supplementary Figure 1B**). Because V2 cells were unsorted, we manually assigned cell types to clusters based on classic markers (**Supplementary Figure 1C-D**). Despite being trained on sparser molecular data from a different version of the Chromium chemistry, classification accuracy remained high, with 80% (3% incorrect, 17% unclassified) of cells correctly labeled with cluster-agnostic type and 95% (3% incorrect, 2% unclassified) with cluster-extended type (**Figure 1D-E**). We also assessed the impact of training on deeply profiled cells in order to classify more sparsely sequenced ones. When trained on V2 cells, Garnett classified V1 cells accurately (83% correct with cluster-agnostic type and 95% correct with cluster-extended) (**Supplementary Figure 1E-G**).

To evaluate Garnett’s ability to catalog cell types in complex solid tissues, we analyzed lung tissue data from two recently reported “molecular atlases” of mouse organs. The Mouse Cell Atlas (MCA)^3^ and Tabula Muris (TM)^2^ projects collected single-cell RNA-seq data using microwell and droplet-based sequencing platforms, respectively. We defined a single hierarchy of cell types expected to be found in the lung based on those studies and compiled marker genes from literature to recognize them in each dataset (all marker files available as **Supplementary Files**, consensus cell type names in **Supplementary Table 1**). Overall, Garnett’s classifications agreed with both the MCA (58% correct, 29% unclassified with cluster-agnostic type; 65% correct, 23% unclassified with cluster-extended type; **Supplementary Figure 2A-B**) and TM (71% correct, 22% unclassified with cluster-agnostic type; 87% correct, 8% unclassified with cluster-extended type; **Supplementary Figure 2C-D**) annotations, which were derived by manual inspection of genes enriched in each cluster. Moreover, a Garnett model trained on the MCA accurately classified the TM cells and vice versa (trained on MCA: 82% correct, 5% unclassified with cluster-agnostic type; 86% correct, 2% unclassified with cluster-extended type; trained on TM: 46% correct, 30% unclassified with cluster-agnostic type; 56% correct, 21% unclassified with cluster-extended type; **Supplementary Figures 2E-H**).

We next sought to evaluate whether Garnett was similarly useful for annotating single-cell chromatin accessibility (scATAC-seq) datasets, which we have generally found to be more challenging to manually annotate than scRNA-seq datasets. We and colleagues recently used regularized, multinomial regression to classify clusters of cells based on chromatin accessibility^8^. We adapted Garnett to classify cells based on scATAC-seq-derived “gene activity scores”, a measure of open chromatin around each gene^13^. Applying it to our recent scATAC-seq atlas of the mouse^8^, Garnett labeled 39% of cells concordantly with our previous assignments (cluster-extended; 22% incorrect; 39% unclassified) (**Supplementary Figure 3**). A caveat is that the marker file was informed by our previous literature-based annotation of the dataset by a related method, but these analyses nonetheless illustrate the potential of Garnett to enable the rapid annotation of not only scRNA-seq but also scATAC-seq datasets.

We next sought to apply Garnett to the task of discriminating all the cell types of a whole animal, focusing on our recent transcriptional atlas of the L2 stage *C. elegans* nematode^14^. We originally assigned broad cell identities to each of 29 major clusters, and then subtyped the neurons using a second level of markers. We defined a cell hierarchy that discriminated the major cell types, as well as subtypes of neurons, using the marker genes from the original study. Of cells that were previously assigned, Garnett labeled 87% of cells concordantly in terms of major cell type (cluster-extended; 8% incorrect; 5% unclassified), with rectum cells being frequently mislabeled as non-seam hypodermis (**Figure 2A-B**, **Supplementary Figures 4-5**). Of the 4,186 neurons assigned subtypes in the original study, 53% were subtyped correctly, and a further 18% were labeled as neurons of unknown subtype (cluster-agnostic; 8% incorrect) (**Figure 2C**). Together, these analyses demonstrate that Garnett can scale to classifying the cell types found in a whole animal.

**Figure 2.**
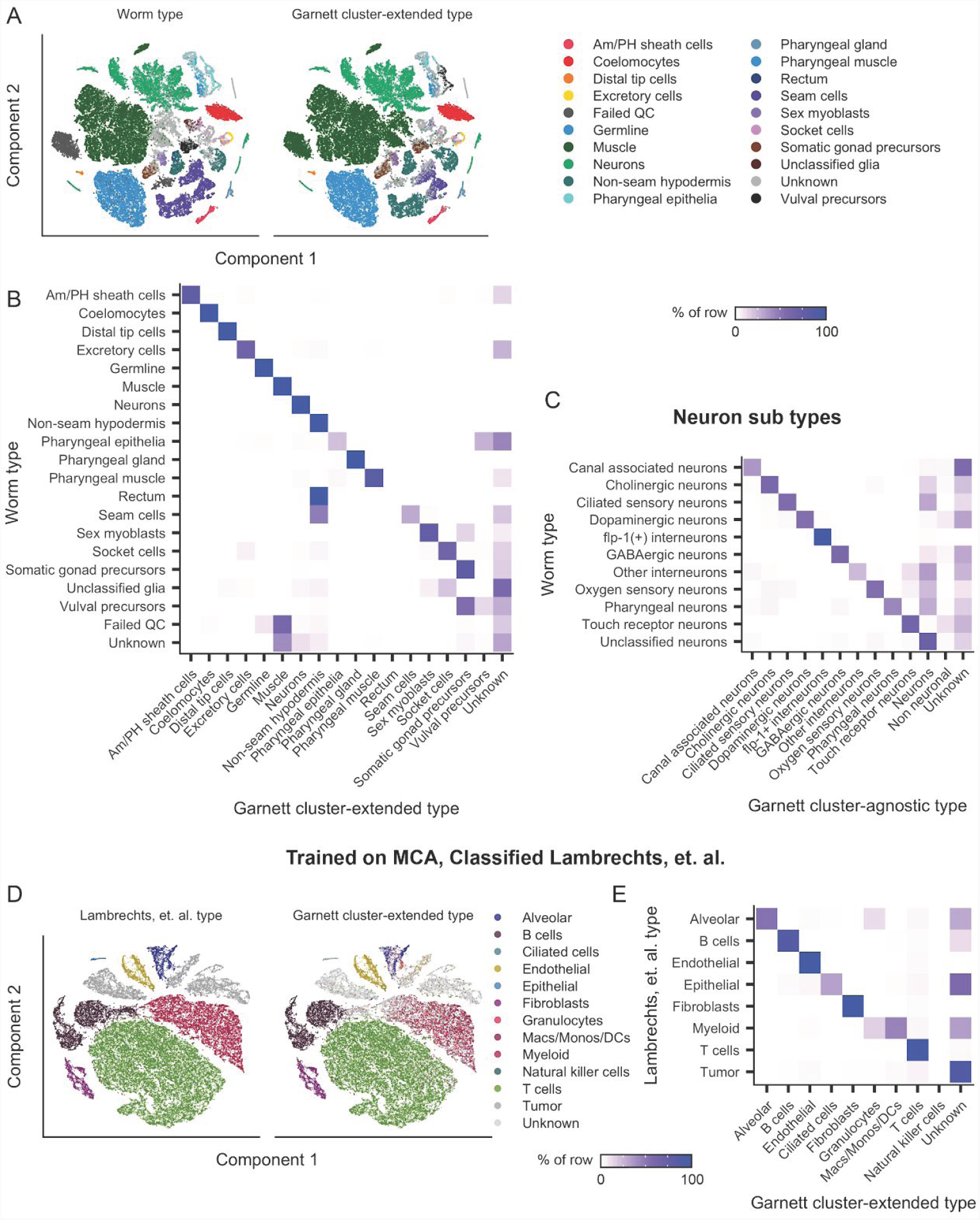
Garnett can discriminate among cell types across a whole animal, across species and between normal and pathological tissue. Garnett classification results for sci-RNA-seq data from whole elegans, published in ref ^14^. **A)** t-SNE plots of the whole worm dataset. First panel is colored by published type from ref ^14^, second panel colored by the major (top level) Garnett cluster-extended classification. Garnett cluster-agnostic type is available in Supplementary Figure 5. **B)** Heatmap comparing the reported cell types versus the Garnett cluster-extended cell types. Color represents the percent of cells of a certain reported type labelled as each type by Garnett. **C)** Heatmap comparing the reported neuron subtypes versus the Garnett cluster-agnostic neuron subtypes. **D)** Garnett cluster-extended results for human lung tumors from ref ^16^ classified based on a Garnett classifier trained on lung cells from the Mouse Cell Atlas. t-SNE plots of the human lung tumor dataset. First panel is colored by published type from ref ^16^, second panel colored by the Garnett cluster-extended classification. **E)** Heatmap comparing the reported cell types versus the Garnett cluster-extended cell types from panel Color represents the percent of cells of a certain reported type labelled as each type by Garnett.

Finally, as tissue--specific patterns of gene expression are largely conserved across vertebrates^15^, we wondered whether Garnett models trained on mouse data could be used to classify human cell types. To evaluate this, we applied the Garnett model trained on the MCA lung dataset to scRNA-seq data from human lung tumors described in ref ^16^ (**Figure 2D-E, Supplementary Figure 6, Supplementary Table 1**). Over 92% of the alveolar, B cells, T cells, epithelial (ciliated) cells, endothelial cells, and fibroblasts were accurately assigned by the Garnett MCA model. Of the 9,756 cells annotated as myeloid^16^, Garnett labeled 44% as monocyte/macrophage/dendritic cell and a further 16% granulocytes, leaving 34% unclassified. 22% of the dataset was labeled “unknown” by Garnett, of which 55% were identified as tumor cells in the original study. As expected, given that they are not represented in the original marker file nor in the MCA lung dataset, 88% of all cells annotated as tumor cells in the original study were labeled as “unknown” by Garnett. These analyses demonstrate that Garnett can operate across species, and is not necessarily confounded by the presence of pathological cell states when trained on normal healthy tissue.

The annotation of cell types based on their molecular signatures is a critical step for the construction of a human cell atlas. It is also increasingly the rate limiting step, as illustrated by recent studies that resorted to labor-intensive, *ad hoc* literature review to achieve this end^2,3,8,14,17,18^. Garnett is an algorithm and accompanying software that automates and standardizes the process of classifying cells based on marker genes. A key point is that the hierarchical marker files on which Garnett is based are interpretable to biologists and explicitly relatable to the existing literature. Furthermore, together with these markup files, Garnett classifiers trained on one dataset are easily shared and applied to new datasets, including across single cell methods and chemistries.

We anticipate the potential for an “ecosystem” of Garnett marker files and pre-trained classifiers that: 1) enable the rapid, automated, reproducible annotation of cell types in any newly generated dataset. 2) minimize redundancy of effort, by allowing for marker gene hierarchies to be easily described, compared, and evaluated. 3) facilitate a systematic framework and shared language for specifying, organizing, and reaching consensus on a catalog of molecularly defined cell types. To these ends, in addition to releasing the Garnett software, we have made the marker files and pre-trained classifiers described in this manuscript available at a wiki-like website that facilitates further community contributions, together with a web-based interface for applying Garnett to user datasets (https://cole-trapnell-lab.github.io/garnett).

## Methods

### Garnett

Garnett is designed to simplify, standardize, and automate the classification of cells by type and subtype. To train a new model with Garnett, the user must specify a cell hierarchy of cell types and subtypes, which may be organized into a tree of arbitrary depth; there is no limit to the number of cell types allowed in the hierarchy. For each cell type and subtype, the user must specify at least one marker gene that is taken as positive evidence that the cell is of that type. Garnett includes a simple language for specifying these marker genes, in order to make the software more accessible to users unfamiliar with statistical regression. Negative marker genes, *i.e.* taken as evidence against a cell being of a given type, can also be specified. In addition, Garnett includes tools for selecting and checking the quality of markers. Garnett uses the marker information provided to select cells that are then used to train a regression-based classifier, similar to the approach taken in ref ^8^. After a classifier is trained, it can be applied to other single cell datasets run on the same or different platforms. Algorithmic details are provided below.

### Constructing marker files

Garnett uses a marker file to allow users to specify cell type definitions. These definitions are then used to choose representative cells from each cell type to use when training the classifier. Full details describing the syntax of the marker file are provided as part of the software package. Briefly, the marker file consists of a series of cell type entries, beginning with a cell type name, followed by lists of expressed markers and metadata. In addition, cell types can be specified to be a subtype of another defined type, *i.e.* hierarchical definitions. Marker files also have the capability to hold literature references for the chosen marker genes that are then included as metadata in the classifier.

Because only markers that are expressed specifically in a given cell type are useful for Garnett classification, we also provide functions for assessing the value of each of the provided marker genes. These functions estimate the number of cells that a given marker nominates for their cell type, the number of cells that become “ambiguously” nominated to multiple cell types in a given level of the hierarchy when the marker is included, and an overall marker score *G*. defined as:

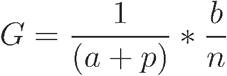

where *a* is the fraction of cells of the cells nominated by the given marker that are made ambiguous by that marker, *p* is a small pseudocount, *b* is the number of cells nominated by the marker, and *n* is the total number of cells nominated for that cell type. In addition to estimating these values, Garnett will plot a diagnostic chart to aid the user in choosing markers (*e.g.* **Supplementary Figure 1A**).

### Training the classifier

Garnett’s first step in training a cell type classifier is to choose representative cells to train on. Let *M* be an *m* by *n* matrix of input gene expression data. First, *M*_*i,j*_ is normalized by size factor (the geometric mean of the total UMIs expressed for each cell *j*) to adjust for read depth, resulting in a normalized *m* by *n* matrix *N*. In addition, the gene IDs of the expression data are converted to Ensembl IDs using correspondence tables from a Bioconductor AnnotationDbi-class ^19^ package. Next, the input marker file is parsed and the gene IDs are also converted to Ensembl IDs as above. Finally, a tree representation of the marker file is constructed, with any designated subtypes placed as children of the parent cell type in the tree. In addition to the tree, a dataset-wide size factor is generated and saved to the tree to allow normalization to new datasets for later classification (see classifying cells section below).

For each parent node in the tree, the following steps are taken: First, cells are scored as “expressed” or “not expressed” for each of the provided markers and an aggregate marker score is derived for each cell type for each cell (details on scoring below). Next, any metadata or hard expression cutoffs are applied to exclude a subset of cells from consideration. Lastly, outgroup samples are chosen (see below). After choosing the training sample, the classifier is trained (see below), and a preliminary classification is made in order to further train downstream nodes.

### Aggregated marker scores

We devised an aggregated marker scoring system to address two challenges of single-cell RNA-seq data for the purposes of identifying representative cell types based on markers. The first challenge when choosing cells is that of differing levels of expression of different markers. If a lowly expressed but specific marker is found in a cell profile, this is better evidence of cell type than a highly expressed and less specific marker. To address this, we use the term frequency-inverse document frequency^20^ (TF-IDF) transformation when generating aggregate marker scores. The TF-IDF transformed matrix is defined by,

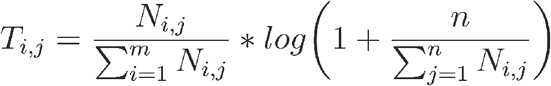

where *N*_*i,j*_ is the *m* by *n* normalized gene expression matrix defined above.

The second challenge we addressed in our aggregate marker score calculation was that highly expressed genes have been known to leak into the transcriptional profiles of other cells. For example, in samples including hepatocytes, albumin transcripts are often found in low copy numbers in non-hepatocyte profiles. To address this, we assign a cutoff above which a gene is considered expressed in that cell. To determine this cutoff we use a heuristic measure defined as

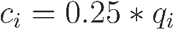

where *c*_*i*_ is the gene cutoff for gene *i* and *q*_*i*_ is the 95th percentile of *T* for gene *i*. Any gene *i* in cell *j* with a value *T*_*i,j*_ below *c*_*i*_ is set to 0 for the purposes of generating aggregated marker scores.

After these transformations, the aggregated marker score is defined by a simple sum of the genes defined as markers in the cell marker file,

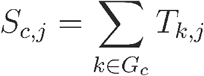

where *S*_*c,j*_ is the aggregated score for cell type *c* and cell *j*, and *G*_*c*_is the list of marker genes for cell type. Cells in the 75th percentile and above for aggregated marker score *S* in only 1 cell type are chosen as good representatives. Any metadata specifications (*e.g.* the requirement that a cell type have come from a particular tissue), provided in the marker file are then used to exclude cells and generate a final training dataset.

### Choosing outgroup cells

When choosing outgroup samples for training, we wanted to make sure that the outgroup set is not dominated by the most abundant cell type. So instead, we cluster a random subset of potential outgroup cells and choose equal numbers of random cells from each cluster to make up the outgroup. Specifically, we first calculate the first 50 principal components using principal components analysis (PCA) as implemented by the irlba^21^ R package. Next, we calculate jaccard coefficients on a k-nearest-neighbors (kNN) graph generated using k = 20. Lastly, we generate clusters using Louvain community detection on the resulting cell-cell map of jaccard coefficients. A random set of cells from each resulting community is then combined to create the outgroup.

### Training with GLMnet

The classifier is trained on the normalized expression matrix *N* for cells chosen as representatives and for all genes expressed in greater than 5% of cells in at least one training set and not expressed in the 90th percentile of TF-IDF transformed expression in all cell types. This last filter prevents ubiquitously expressed genes from being chosen as features. The classifier is trained using genes as features and cells as observations with a grouped multinomial elastic net regularized (alpha = 0.3) generalized linear model using the package GLMnet^22^ in R. Observations are weighted by the geometric mean of the counts in each of the training groups. The GLMnet regularization parameter λ is chosen using 3-fold cross validation. Genes provided in the marker files are required to be included in the model not regularized.

### Classifying cells

Because we wished to be able to use pre-trained classifiers to classify cells across datasets and platforms, we include a dataset size factor *D* for the training data with the classifier object. *D* is the geometric mean of the total read counts per cell divided by the median number of genes expressed above zero per cell. Formally, *D* is defined by

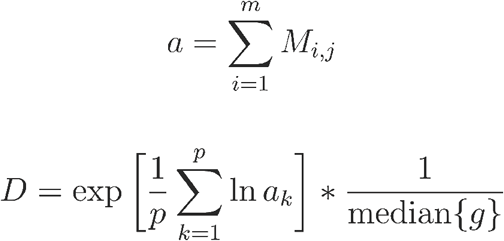

where *g* is the number of genes expressed above zero per cell. When applying an existing classifier to a new dataset, we can then transform the new expression data, an *m*′ by *n*′ matrix *M*′, to the scale of the training data using *D*,

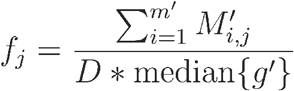

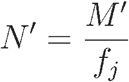

where *g*′is the number of genes expressed above zero per cell in the new data.

After normalization, gene IDs for the new dataset are also converted to Ensembl IDs. At each internal node in the classifier, the multinomial model for that node is applied to the data, the output probabilities of each class are normalized by dividing by the maximum probability for each cell, and the ratio of the top scoring cell type to the second best scoring cell type is calculated. If this odds ratio is greater than the user-specified rank probability ratio (in this paper and by default, we use 1.5), the top type is assigned, otherwise the cell type is set to “Unknown”. Optionally, Garnett will add a second set of classifications which classify an entire cluster of cells if: greater than 90% of assigned cells within a cluster are the same type and greater than 5% of all cells in the cluster are classified (not “Unknown”), and greater than 5 cells in the cluster are classified. Cluster labels can be provided by the user or generated by Garnett using Louvain community detection in the top 50 principal components of the expression matrix.

### 10x Genomics Peripheral Blood Mononuclear Cells (PBMCs)

10x PBMC datasets from both version 1 (V1) and version 2 (V2) chemistry were downloaded from the 10x Genomics website. The V1 cells are a combination of each of the pure cell type populations isolated by 10x Genomics using FACS sorting (CD14+ Monocytes, CD19+ B cells, CD34+ cells, CD4+ Helper T cells, CD4+/CD25+ Regulatory T cells, CD4+/CD45RA+/CD25- Naive T cells, CD4+/CD45RO+ Memory T cells, CD56+ Natural killer cells, CD8+ Cytotoxic T cells and CD8+/CD45RA+ Naive cytotoxic T cells) preprocessed using CellRanger 1.1.0 and published in Zheng et. al. The V2 cells are the V2 chemistry distributed demonstration dataset labelled “8k PBMCs from a healthy donor”, preprocessed using CellRanger 2.1.0. Markers for PBMCs were those often cited in the literature. Using Garnett’s marker scoring system, we excluded the markers with high ambiguity (>0.5). The final PBMC marker file used is available as Supplementary Data File 1. Garnett classification for V1 and V2 was run using default parameter values defined in the preceding sections.

### Tabula muris and mouse cell atlas MCA lung analysis

The Tabula muris FACS dataset from ref ^2^ was downloaded from their figshare website. The MCA dataset from ref ^3^ was downloaded from their figshare website. For the purposes of this analysis, only data derived from lung tissue from both datasets were used. To facilitate comparisons between each of the lung datasets used, a set of consensus cell type names were used as described in Table 1. The marker file used is available as Supplementary Data File 2. Garnett classification was run using default parameter values for both datasets.

### sci-ATAC-seq analysis

The sci-ATAC-seq data was downloaded from the website associated with ref ^8^. The input to Garnett was the previously calculated Cicero gene activity scores presented in the original publication. The final marker file used is available as Supplementary Data File 3. Garnett classification was run using default parameter values.

### Worm analysis

The worm data was downloaded from the website associated with ref ^14^. Markers were those used by the original publication to identify cell types. Using Garnett’s marker scoring system, we excluded the markers with high ambiguity (Supplementary Figure 4). The final marker file used is available as Supplementary Data File 4. Garnett classification was run using default parameter values.

### Human lung tumor analysis

The human lung tumor data was downloaded from the ArrayExpress database entry associated with ref ^16^. Because expression data were log-transformed, we first exponentiated the expression data before classification. To allow for cross-species classification, we first converted the human expression data to mouse gene labels by creating a correspondence table using the biomaRt hsapiens_gene_ensembl and mmusculus_gene_ensembl databases. Only unique rows (one-to-one correspondences) were used. Ultimately 15,336 of the original 22,180 human genes could be converted to mouse labels including 89 percent of the genes in the MCA classifier with non-zero coefficients. The final marker file used is available as Supplementary Data File 5. Garnett classification was run using default parameter values.

## Data Availability

No new data was generated for this study. All data used in this study is publicly available.

## Software Availability

Garnett is an R package available through github.

## Acknowledgements

We gratefully acknowledge Stephen Tapscott, William Noble, and Daniela Witten as well as members of the Shendure and Trapnell labs, particularly Andrew Hill, for their advice. Zachary Pliner named the software. This work was supported by the following funding: NIH grant U54DK107979 to JS and CT; NIH grant DP2HD088158, RC2DK114777 and R01HL118342 to CT; NIH grants DP1HG007811 and R01HG006283 to JS; and the Paul G. Allen Frontiers Group to JS and CT. JS is an Investigator of the Howard Hughes Medical Institute. CT is partly supported by an Alfred P. Sloan Foundation Research Fellowship. HAP was supported by the National Science Foundation Graduate Research Fellowship under Grant No. (DGE-1256082).

---

**Supplementary Figure 1.**
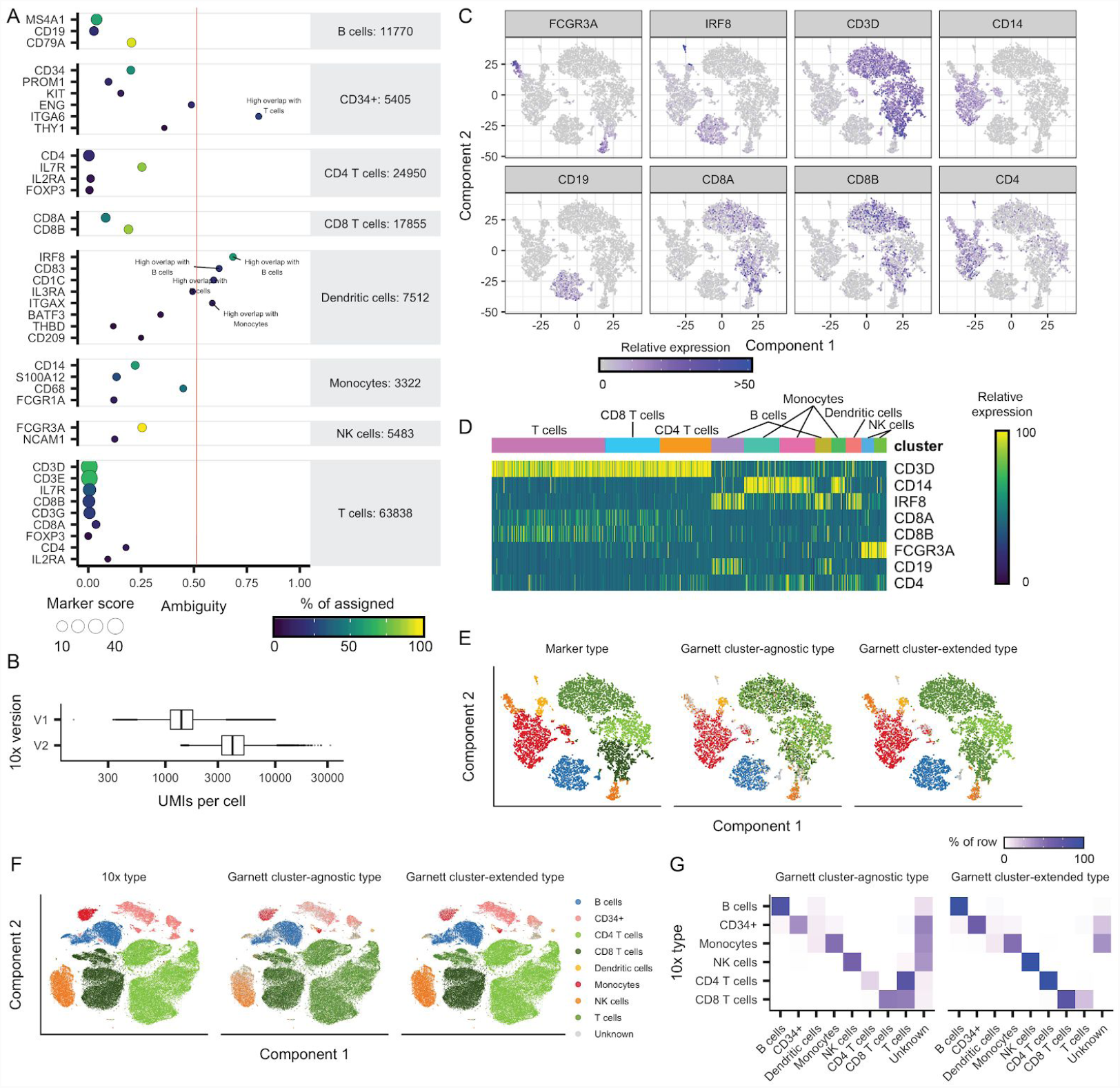
Garnett accurately classifies peripheral blood mononuclear cells. A) PBMC marker quality plot. X-axis corresponds to the ambiguity score, defined as the ratio of the number of ambiguous cells when the marker is included over the number of cells in which the marker is expressed. Color represents the percent of nominated cells for that cell type that were nominated by that marker, and the number next to the cell type names is the total number of nominated cells in that cell type. Markers with an ambiguity score greater than 0.5 (indicated by the red line) were excluded from the marker file. **B)** Boxplots of the number of unique molecular indexes (UMIs) per cell in 10x Genomics version 1 (V1) PBMC dataset versus version 2 (V2) (Boxplot elements: center line, median; box limits, upper and lower quartiles; whiskers, 1.5x interquartile range; points, outliers). **C)** t-SNE plots of 10x Genomics V2 PBMC dataset. Color represents the relative expression of marker genes for each expected cell type (FCGR3A: NK cells, IRF9: Dendritic cells, CD3D: T cells, CD14: Monocytes, CD19: B cells, CD8A and CD8B: CD8 T cells, CD4: CD4 T cells). **D)** Correspondence between markers of interest and cell clusters in 10x Genomics V2 PBMC dataset with manually assigned cell type labels. Heatmap of relative expression, rows are marker genes and columns are cells sorted by t-SNE cluster assignment. **E)** t-SNE plot of Garnett cluster-agnostic and cluster-extended type assignments for 10x Genomics V2 PBMCs, also trained on V2. **F)** t-SNE plot of Garnett cluster-agnostic and cluster-extended type assignments for 10x Genomics V1 PBMCs, trained on V2. **G)** Correspondence of Garnett cluster-agnostic and cluster-extended type assignments with FACS assignments for data from (F). Color represents the percent of cells of a certain FACS type labeled each type by Garnett.

**Supplementary Figure 2.**
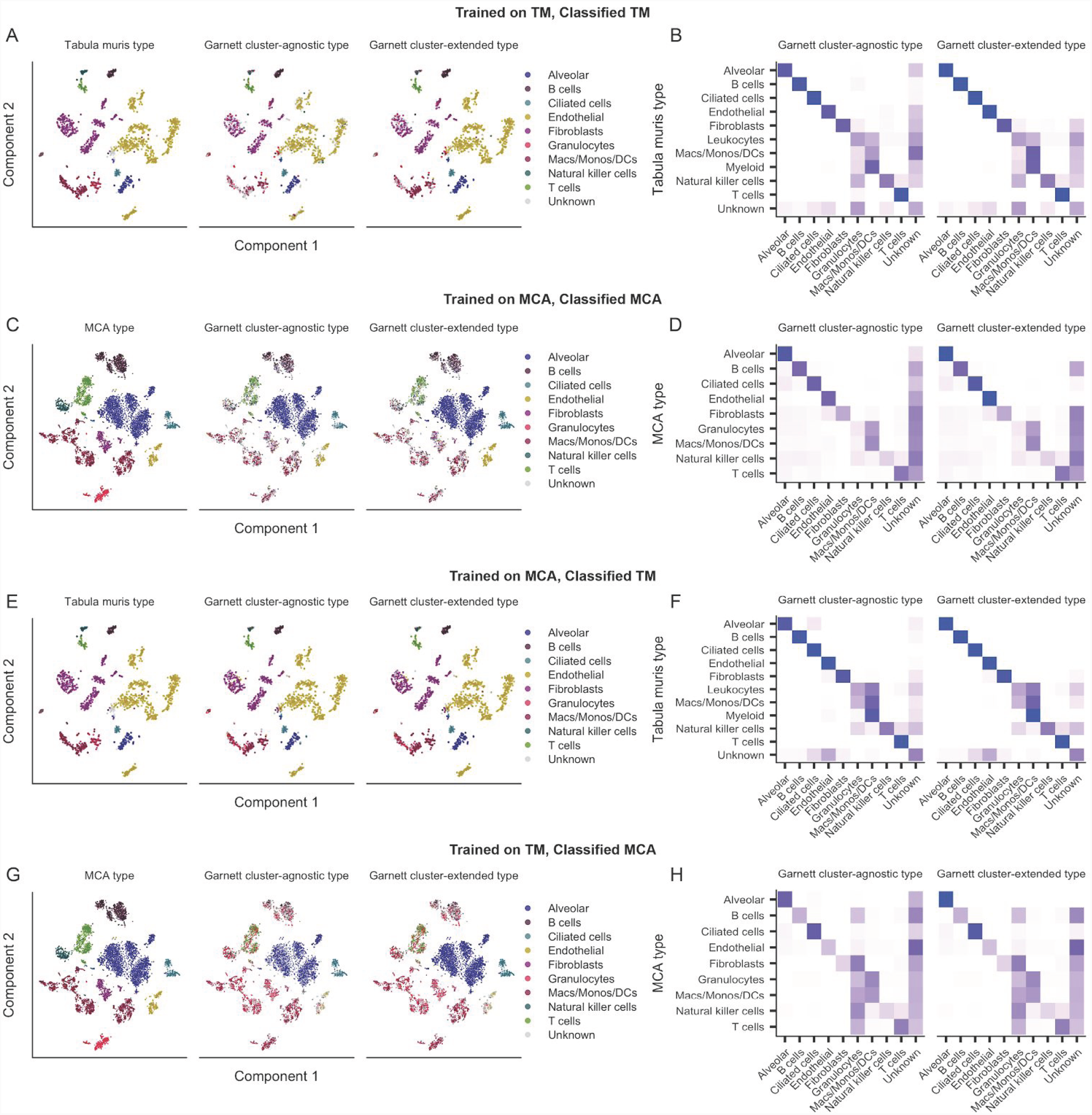
Garnett accurately classifies lung cell types from recent mouse cell atlases. Panels **A**, **C**, **E**, and **G** are t-SNE plots of Tabula Muris (TM)^2^ and Mouse Cell Atlas (MCA)^3^ lung subsets colored by reported cell type versus Garnett cluster-agnostic and cluster-extended types. Panels **B**, **D**, **F** and **H** are heatmaps comparing the reported cell types (rows) versus the Garnett cluster-agnostic and cluster-extended types (columns). Color represents the percent of cells of a certain reported type labelled as each type by Garnett. Panels B, D, F and H correspond with A, C, E, and G respectively. Panels A and B are TM data, and were classified using the TM-trained classifier. Panels C and D are MCA data, and were classified using the MCA-trained classifier. Panels E and F are TM data, and were classified using the MCA-trained classifier. Panels G and H are MCA data, and were classified using the TM-trained classifier.

**Supplementary Figure 3.**
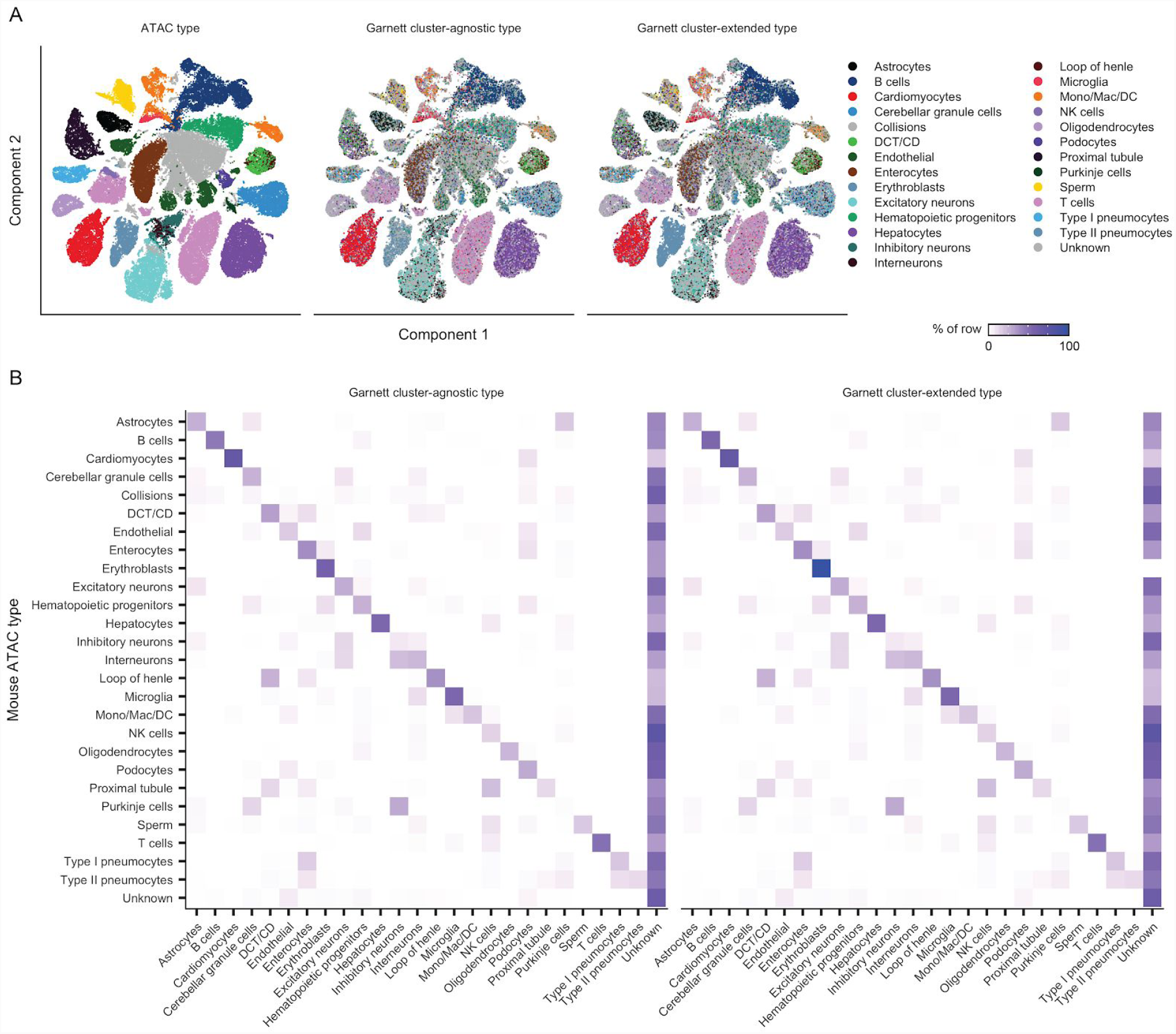
Garnett can classify cells from single-cell chromatin accessibility datasets. A) t-SNE plot of the Cusanovich *et al.^8^* mouse single-cell ATAC-seq atlas. Garnett used publicly available Cicero^13^ gene activity scores in place of expression data to classify cell types. The first panel is colored by Cusanovich *et al.^8^* manually assigned cell type labels. The second and third panel are colored by the Garnett cluster-agnostic and cluster-extended types respectively. **B)** Heatmaps comparing the reported cell types versus the Garnett cluster-agnostic and cluster-extended types. Color represents the percent of cells of a certain reported type labelled as each type by Garnett.

**Supplementary Figure 4.**
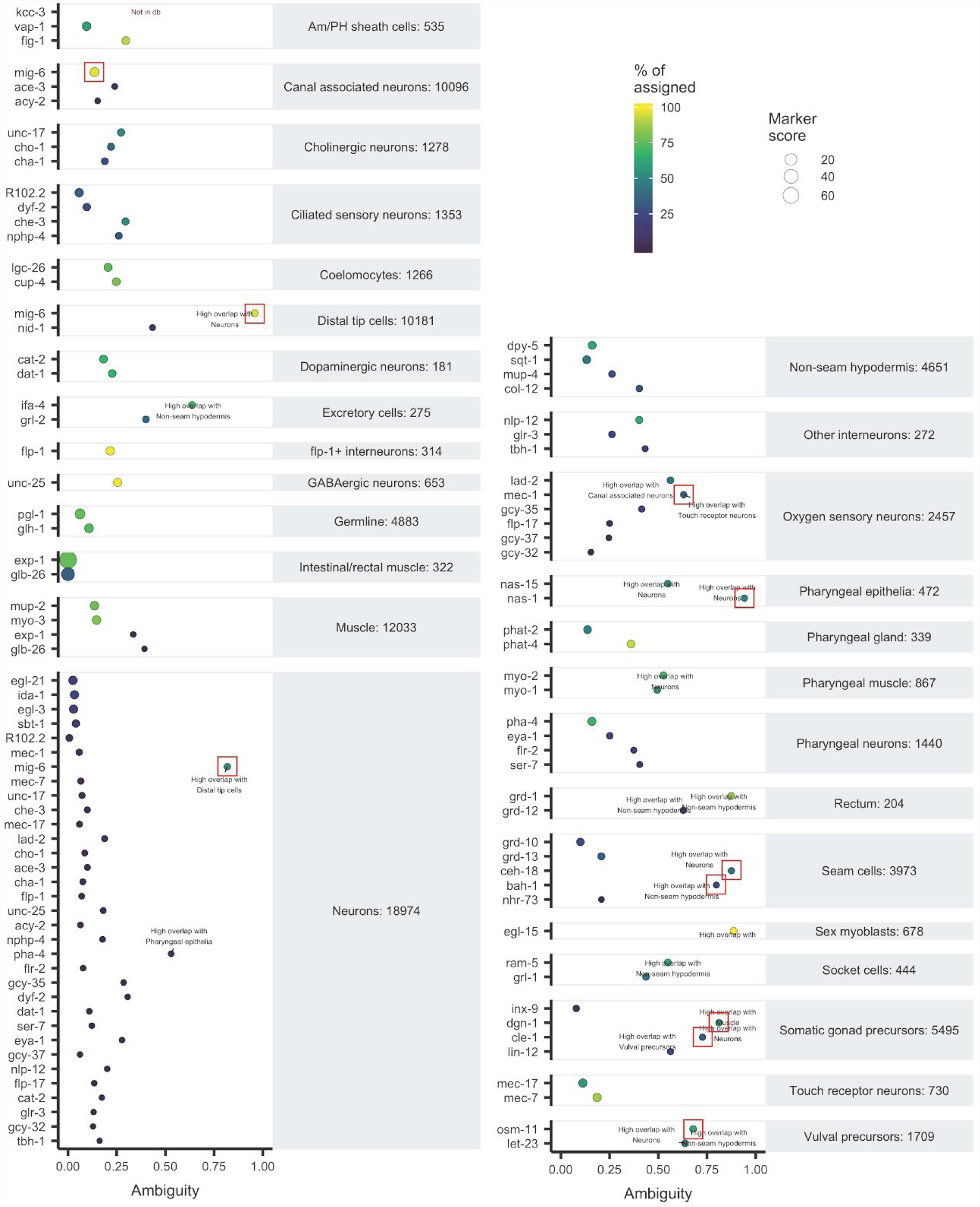
Marker quality chart for *C. elegans*. X-axis represents the ambiguity score, defined as the ratio of number of ambiguous cells when the marker is included over the number of cells the marker is expressed in. Color represents the percent of nominated cells for that cell type that were nominated by that marker, and the number next to the cell type names is the total number of nominated cells in that cell type. Markers were initially chosen directly from ref ^14^. Markers excluded because of high ambiguity are marked with red boxes.

**Supplementary Figure 5.**
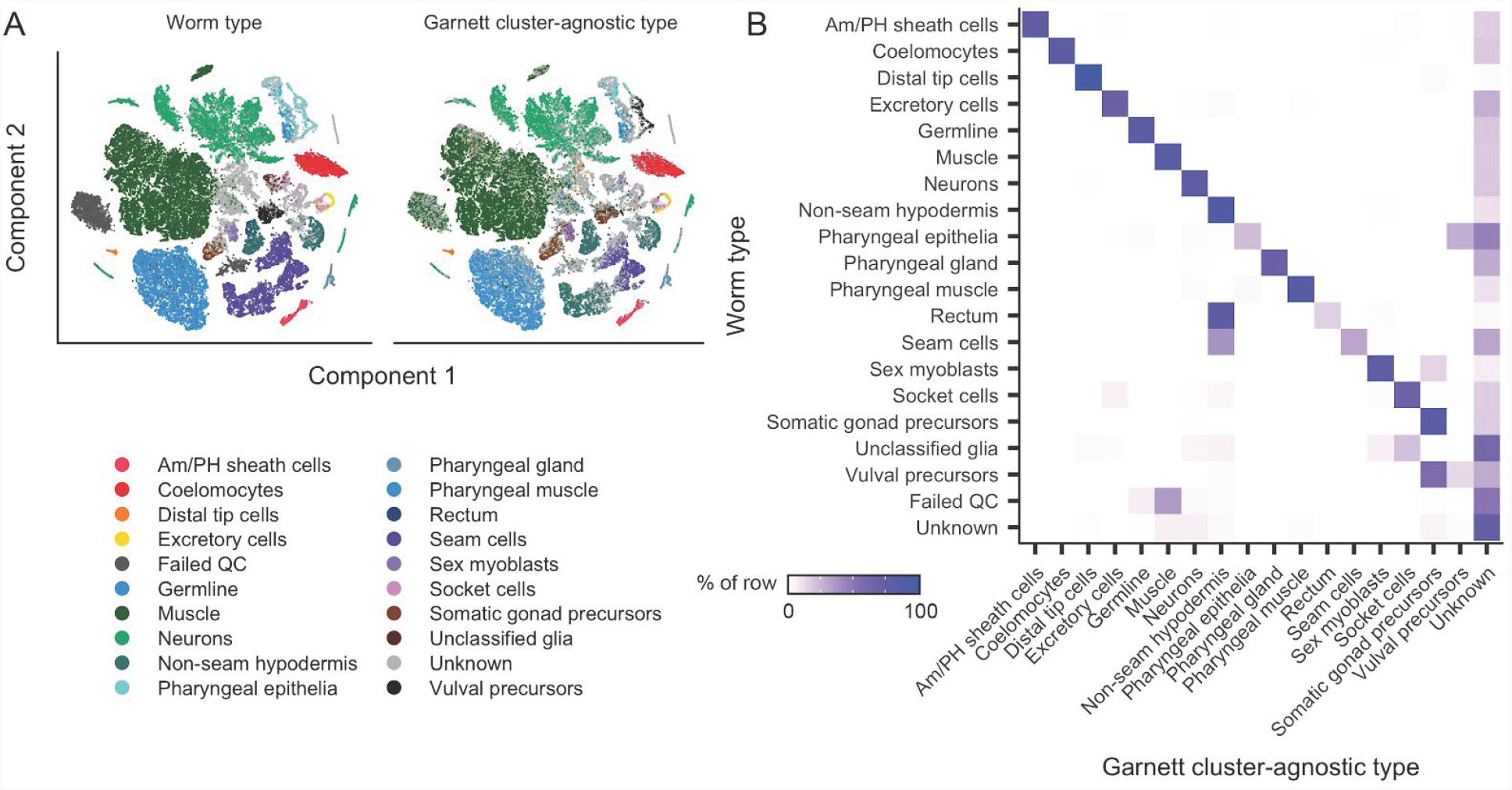
Garnett classification results for sci-RNA-seq data from whole L2 stage *C. elegans*. A) t-SNE plots of the whole worm dataset^14^. First panel is colored by published type from ref^14^, second panel colored by the major (top level) Garnett cluster-agnostic classification. **B)** Heatmap comparing the reported cell types versus (rows) the Garnett cluster-agnostic cell type (columns). Color represents the percent of cells of a certain reported type labelled as each type by Garnett.

**Supplementary Figure 6.**
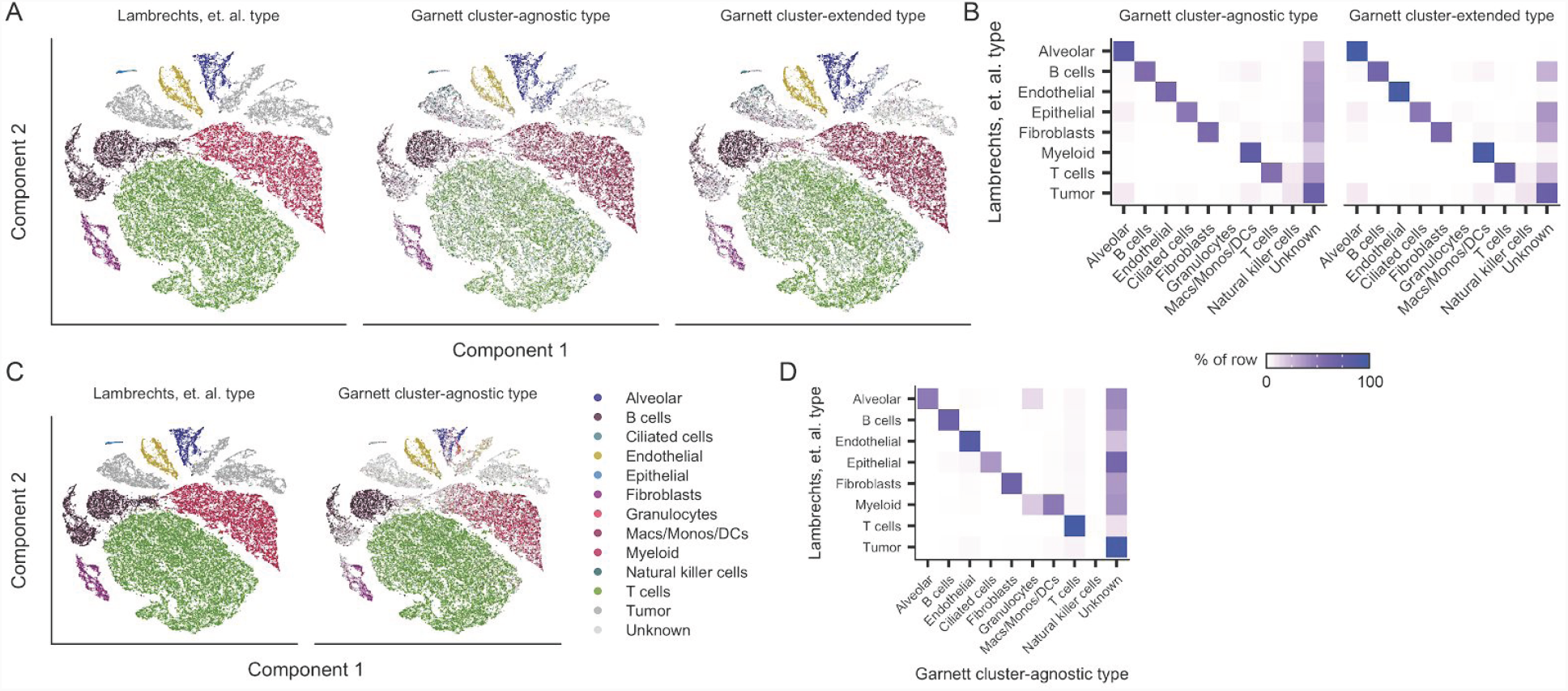
Garnett classification of single-cell RNA-seq data from lung tumors. A) t-SNE plots of lung tumor scRNA-seq dataset^16^. First panel is colored by published type from ref ^16^, second panel colored by Garnett cluster-agnostic cell type, and third panel colored by Garnett cluster-extended cell type, based on a model trained using this same dataset. **B)** Heatmaps comparing the reported cell types versus the Garnett cluster-agnostic and cluster-extended cell types from panel A. Color represents the percent of cells of a certain reported type labelled as each type by Garnett. **C)** Garnett cluster-agnostic results for human lung tumors from ref ^16^ classified based on a Garnett classifier trained on lung cells from the Mouse Cell Atlas. t-SNE plots of the human lung tumor dataset. First panel is colored by published type from ref ^16^; second panel colored by the Garnett cluster-agnostic classification. **D)** Heatmap comparing the reported cell types versus the Garnett cluster-agnostic cell types from panel C. Color represents the percent of cells of a certain reported type labelled as each type by Garnett.

**Supplementary Table 1.** Consensus cell types for lung datasets.

| Consensus type | Tabula muris type | MCA type | Lambrechts type |
| --- | --- | --- | --- |
| Alveolar | epithelial cell of lung | AT2 Cell<br>AT1 Cell<br>Alveolar bipotent progenitor | Alveolar |
| B cells | B cell | Ig-producing B cell<br>B Cell | B cells |
| Ciliated cells | ciliated columnar cell of tracheobronchial tree | Ciliated cell<br>Clara Cell<br>Dividing cells | Epithelial |
| Endothelial | lung endothelial cell | Endothelial cell_Tmem100 high<br>Endothelial cells_Vwf high<br>Endothelial cell_Kdr high | Endothelial |
| Fibroblasts | stromal cell | Stromal cell_Inmt high<br>Stromal cell_Dcn high<br>Stromal cell_Acta2 high | Fibroblasts |
| Granulocytes |  | Neutrophil granulocyte<br>Eosinophil granulocyte<br>Basophil |  |
| Leukocytes | leukocyte |  |  |
| Macs/Monos/DCs | classical monocyte<br>monocyte | Conventional dendritic cell_H2-M2 high<br>Dendritic cell_Naaa high<br>Dividing dendritic cells<br>Conventional dendritic cell_Tubb5 high<br>Conventional dendritic cell_Gngt2 high<br>Plasmacytoid dendritic cell<br>Conventional dendritic cell_Mgl2 high<br>Monocyte progenitor cell<br>Interstitial macrophage<br>Alveolar macrophage_Ear2 high<br>Alveolar macrophage_Pclaf high |  |
| Myeloid | myeloid cell |  | Myeloid |
| Natural killer cells | natural killer cell | NK Cell |  |
| T cells | T cell | T Cell_Cd8b1 high<br>Dividing T cells<br>Nuocyte | T cells |
| Tumor |  |  | Tumor |

